# BioPAX-Explorer: a Python Object-Oriented framework for overcoming the complexity of querying biological networks

**DOI:** 10.1101/2024.09.18.613626

**Authors:** François Moreews, Jean-Baptiste Bougaud, Emmanuelle Becker, Florence Gondret, Olivier Dameron

**Affiliations:** Univ Rennes, Inria, CNRS, IRISA, France; Pegase, Inrae, Institut Agro, 35590 Saint-Gilles, France

## Abstract

**Motivation:** Biological Pathway Exchange (BioPAX) is a standard language, represented in OWL, that aims to enable the integration, exchange, visualization and analysis of biological pathway data. While public databanks increasingly provide datasets in BioPAX format, their use remains below potential. Users may encounter challenges in harnessing the data due to the BioPAX intricately detailed underlying model. Moreover, extracting data demands specific technical skills, posing a barrier for many potential users.

**Results:** To address these obstacles, we developped BioPAX-Explorer. This toolis designed to facilitate the adoption and usage of BioPAX for extracting data or build algorithms and models, within the Python community. BioPAX-Explorer is a Python package that provides an object-oriented data model automatically generated from the BioPAX OWL specification. Moreover, it offers expressive query capabilities that shield users from BioPAX inner complexity. BioPAX-Explorer supports dataset building features, validation facilities and pre-build queries. It simplifies the extraction and processing of data from BioPAX sources by automatically generating SPARQL queries. BioPAX-Explorer also offers a user-friendly interface for Python users, allowing exhaustive exploration of large datasets through features such as memory-efficient query execution, entity-oriented queries without the need for SPARQL knowledge. It also allows to learn and reuse complex SPARQL queries for biological network analysis. Additionally, BioPAX-Explorer can accelerate the development of Python-based network analysis software, since it generates graph data structures from BioPAX queries and facilitates the creation of transparent, reproducible workflows based on the BioPAX OWL standard.

**Availability and implementation:** BioPAX-Explorer is freely available. We provide the source code, documentation, installation instructions and a Jupyter notebook with tutorial at https://fjrmoreews.github.io/biopax-explorer/

## 1. INTRODUCTION

The diverse and complex nature of biological networks, which encompass molecular interactions between entities and reconstruction processus, necessitates the development of standardized formats and software frameworks to integrate data from various complementary sources.

Addressing this challenge is crucial for understanding of biological processes and facilitating collaborative efforts in the analysis and interpretation of comprehensive network datasets. Biological Pathway Exchange (BioPAX) plays a pivotal role in standardizing and ensuring the interoperability of biological network data (Demir *et al*., 2010). Indeed, BioPAX serves as a standardized language with the primary goal of facilitating the exchange, integration, visualization, and analysis of biological pathway data. Represented in RDF and defined in OWL, BioPAX has become a common format for sharing datasets in public data banks. Despite the increasing availability of BioPAX files, potential users often face challenges in leveraging the valuable data due to the complexity of the underlying model. As a matter of fact, the complete BioPAX Level 3 data model consists of over 60 type of entities (classes) with complex and intertwined relationships, and the textual documentation covers more than 160 pages. Moreover, exploiting BioPAX requires specialized technical skills for data extraction.

To facilitate the adoption of BioPAX, different tools have been proposed. Paxtools (Demir *et al*., 2013) is a widely adopted implementation with comprehensive functionality for manipulating BioPAX datasets, offering high expressiveness in pattern-based queries (Babur *et al*., 2014). However, it has drawbacks such as in-memory dataset handling requirements, non-standard query system that can complicate the understanding of query patterns, and the requirement for queries to be implemented in Java, which may limit its adoption within the bioinformatics community. PaxtoolsR (Luna *et al*., 2016) serves as a straightforward R wrapper facilitating the seamless integration of the Java-based Paxtools tool into the R environment. It acts as an interface that allows communication between R and Paxtools, enabling R users to leverage Paxtools functionalities without the need for direct Java programming.

More recently, PyBioPAX (Gyori and Hoyt, 2022) offers comprehensive functionality for processing, manipulating, and interacting with BioPAX models in Python. It supports model serialization and implements BioPAX OWL semantics. It is integrated with web services for knowledge extraction, but is currently limited to the Pathway Commons database. PyBioPAX allows to represent BioPAX entities as instances of Python classes but does not implement any expressive query system. Data extraction and filtering are carried out in memory after the entity instances have been created, leading to large memory footprint.

Tools like BioCypher (Lobentanzer *et al*., 2023) are also available to prototype a knowledge graph data infrastructure. BioCypher employs an object-oriented paradigm, and a graph databases (Neo4j) to facilitate the integration of data into a knowledge graph. However, it does not ensure interoperability between datasets that may be shared by different teams, because it is not built on standards.

Therefore, there is a lack of tools designed to streamline the querying of all BioPAX datasets, specifically tailored for Python users and compliant with established standards. BioPAX-Explorer was designed to address these requirements and offer a tool suited to the needs of bioinformaticians familiar with Python. The primary function of BioPAX-Explorer is to query BioPAX datasets, while guaranteeing compatibility with Semantic Web standards to ensure a high degree of interoperability and openness.

We first delineate the primary goals of BioPAX-Explorer based on user requirements, providing insight into the design choices that have been made. We then describe the technologies and techniques employed for the underlying framework. Then, we list the main functionalities implemented in BioPAX-Explorer and provide showcases to illustrate different applications. Future directions and refinement opportunities are also indicated for contributing to the advancement of interoperability solutions in the exploration of biological networks.

## 2. Approach

We aimed to simplify the use of BioPAX for data extraction and algorithm design, targeting Python users, who are prevalent in bioinformatics. Figure 1 summarizes the expected features of BioPAX-Explorer.

**Fig 1.**
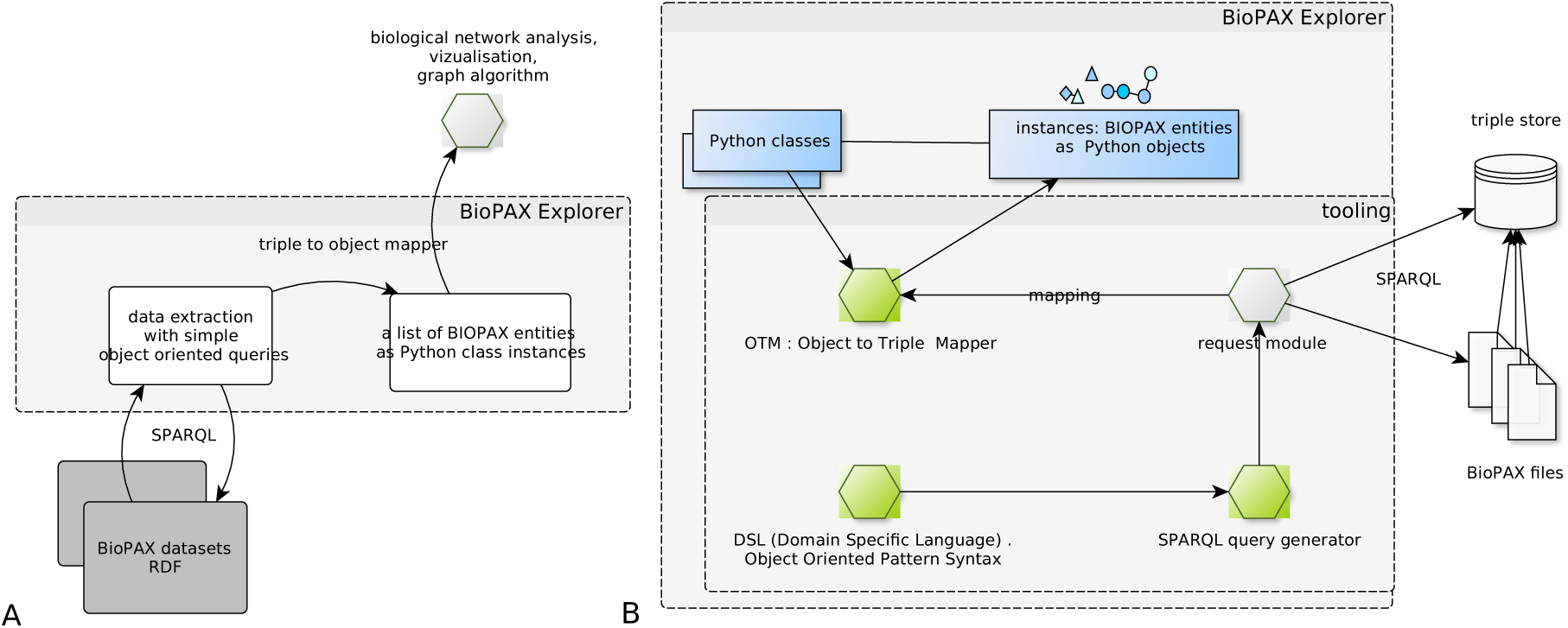
Main functionalities of BioPAX-Explorer (A) and software component architecture (B). We summarized here the expected features of our tool to simplify querying BioPAX datasets. The desirable functionalities of BioPAX-Explorer should include an ability to simplify BioPAX dataset querying through the creation of Python-based object-oriented queries. These queries must be automatically translated into SPARQL, the standard language for RDF data extraction. Subsequently, they can be executed using an engine, with the results seamlessly transformed into Python class instances corresponding to BioPAX model entities. Either files or a triple store can be used as a data source.

Our approach is based on leveraging the web ontology language OWL and the Resource Description Framework RDF, technologies used to define BioPAX. This strategy eliminates the needs for recoding model specifications and parsing datasets, two tasks that can introduce errors. OWL is a Semantic Web language that allows the definition and representation of ontologies, enabling automated and knowledge-based processing of information. Hence, it is possible to use code generation to derive a data model from OWL. RDF is a serialization format for Semantic Web data, providing a standardized way to represent and exchange data in a machine-readable format. Considering that BioPAX datasets are defined in RDF, it is adapted to SPARQL (Protocol and RDF Query Language) for querying and retrieving information stored in RDF format. SPARQL is used in the Semantic Web for flexible and expressive data retrieval, but requires specific technical expertise, and demands a thorough understanding of the data model being manipulated.

To overcome these limitations, we provide a tool that generates SPARQL queries for the user. However, this raises the question of what type of programming method has to be proposed to users for defining queries. We opted to provide a simplified programming interface (API) for defining search queries, that are automatically converted to SPARQL upon execution.

For this purpose, we created an API allowing users to define queries in the form of search patterns, following an object-oriented paradigm that is easy to grasp as it is close to the BioPAX data model. We also sought to ensure that the query results will be instances of model entities rather than RDF triples, simplifying data manipulation for the users. The object-oriented model is familiar for developers because it mirrors the real-world organization of entities, promoting encapsulation, modularity, and abstraction.

We describe here the concepts and methods used.

### 2.1 Triple to Object Mapper

RDF triples consist of subject-predicate-object representations. Within the field of Linked Data and RDF, a “triple to object mapper” refers to a component, that specifies the mapping between an object and its representation as RDF triples. These mappers help to bridge the gap between RDF data and the programming language’s object-oriented paradigm (Quasthoff and Meinel, 2012). Thus, RDF triples are represented as objects, which can have properties, methods, and relationships in accordance with the programming language’s constructs. Figure 2 displays a graphical representation of a BioPAXobject in the form of RDF data (A) and a possible corresponding class (B) in the classic object-oriented modelling paradigm.

**Fig 2.**
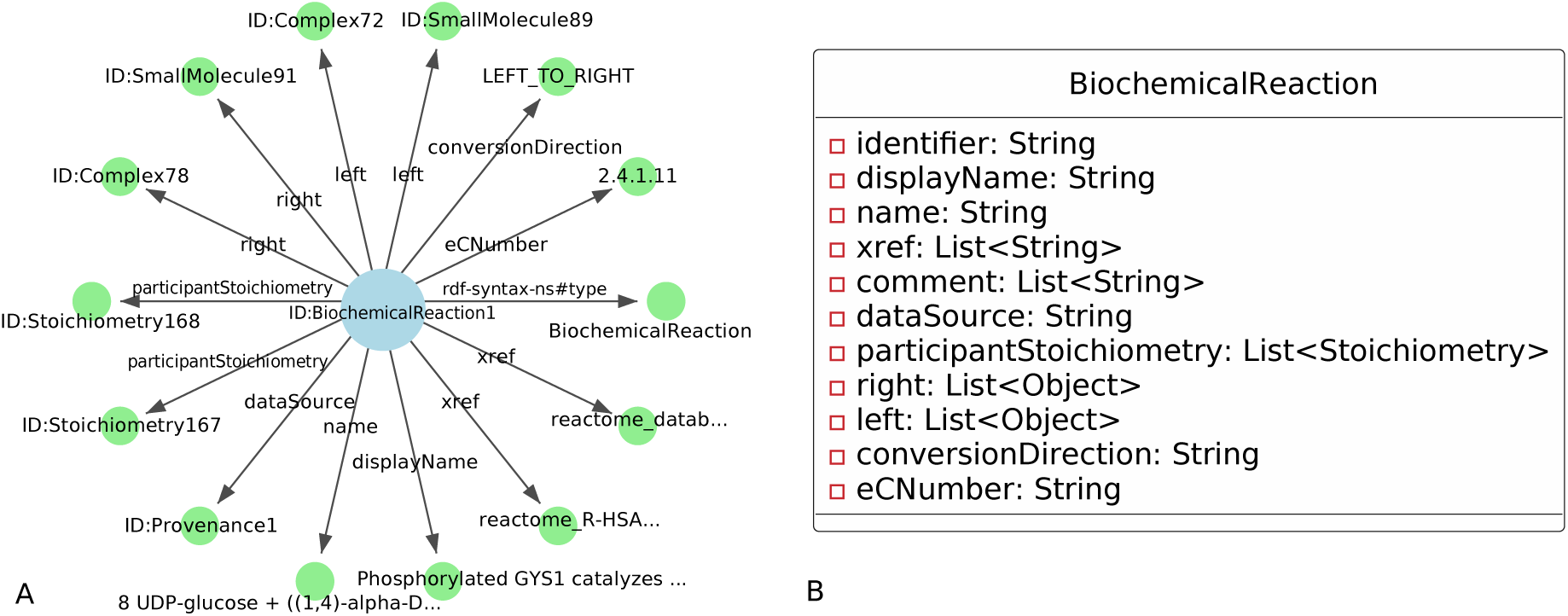
Graphical representation of an instance of the class BiochemicalReaction that complies to the BioPAX specification (A). The source data consists of RDF triples, represented here in the form of a star whose centre is the object identifier or URI (Uniform Resource Identifier). The underlying class (B) acts as a general data structure shared by all BiochemicalReaction instances (objects). The Triple to Object Mapper enables the automated conversion of RDF data into such structures and vice versa, allowing easy manipulation of the data in a programming language such as Python.

BioPAX-Explorer offers this mapping functionality, allowing the automated translation of triples into objects and vice versa, as described in Figure 1.

### 2.2 Compliance with the BioPAX standard

To ensure a robust compatibility with the BioPAX standard, we use a code generation technique to create the Python model classes from the OWL specification. For that, we first parse the OWL BioPAX specification and then, generate Python classes representing BioPAX entities. While this method may require time, it ensures a high level of compatibility and quality, tailored to this extensive model.

Figure 3 illustrates this automated process, where the OWL specification is translated into a set of corresponding Python classes.

**Fig 3.**
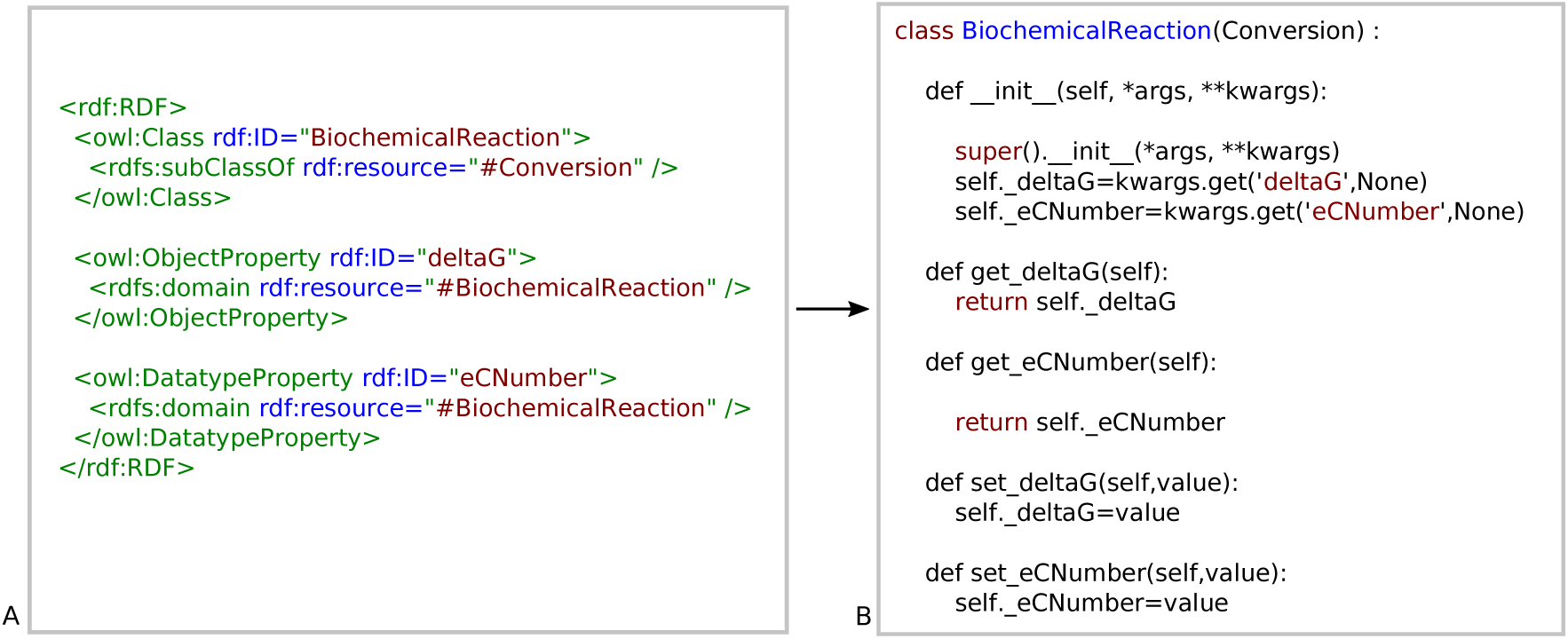
Compliance to the BioPAX standard is ensured through a code generator that automatically translates the OWL specification (A) into Python code (B). A distinct “.py” file is produced for each BioPAX class. Here, we present the class BiochemicalReaction for illustration. For enhanced clarity, only a partial view of the OWL source and the corresponding generated Pythoncode is shown, but detailed aspects of data typing is managed too, like list and range. This process was implemented during the BioPAX-Explorer development phase.

### 2.3. Designing a intuitive query language

A domain-specific language (DSL) for querying defines queries and patterns specific to the structures and relationships within that applicative domain. This tailored language allows users to proficiently retrieve and analyze pertinent data, featuring syntax designed for the nuances of the domain’s representations. This key component of BioPAX-Explorer transforms user-defined patterns into SPARQL query graphs. It is responsible for translating high-level queries into executable SPARQL queries. This functionality empowers users to express complex queries in a concise and intuitive manner. The DSL BioPAX pattern language is integrated in BioPAX-Explorer as an API.

## 3. Materials and methods

The software package of BioPAX-Explorer is implemented in Python 3, ensuring compatibility with version 3.8 and higher versions. It leverages an OWL-based code generation module, by utilizing custom codes and the OWLReady2 (Lamy, 2017)^1^ package which facilitates the efficient manipulation of OWL ontologies. For graph management tasks, the package employs NetworkX (Hagberg *et al*., 2008) as its primary graph manipulation library. Additionally, it provides optional support for Graph-tool (Peixoto, 2017), which is particularly useful for handling large graphs efficiently in memory. This dual-library approach ensures flexibility and scalability, to adapt to various graph sizes and computational resources.

Execution of SPARQL queries is implemented through the RDFLib ^2^ package, a widely-used library for working with RDF data. RDFLib provides robust functionality for interacting with RDF data sources, including file-based storage and triple stores. This enables the package to query and manipulate RDF data, that can be stored locally or remotely.

A custom code generator, independently developed, produces the Python classes corresponding to the BioPAX model. This utility tool operates by ingesting the OWL specification file, which serves as the authoritative source defining the BioPAX format specification. Upon receiving the OWL specification file as input, the code generator employs a parsing mechanism to extract relevant information about classes, properties, and relationships specified within the ontology. Subsequently, it utilizes this parsed data to generate Python class definitions that accurately reflect the structure and semantics of the BioPAX model. The code generator employs Jinja2 ^3^ to manage code templates. These templates serve as the blueprint for generating Python class files, incorporating placeholders for inserted data extracted from the OWL BioPAX specification. This modular approach permitted to easily modify or extend the generated code by adjusting the underlying templates.

Once code generation is complete, the output comprises a collection of Python files containing the generated model classes. To ensure the correctness and functionality of these classes, automated tests where conducted as part of the code generation process. These tests encompass scenarios such as instantiating classes from BioPAX files and validating the integrity of generated BioPAX files derived from class instances. During testing, instances of the generated Python classes are programmatically created from sample BioPAX files, mimicking real-world usage scenarios. Subsequently, these instances undergo validation to ensure adherence to the BioPAX specification and proper handling of data and relationships.

Additionally, the generation of BioPAX files from instantiated class objects is scrutinized to verify the accuracy and completeness of the serialization process.

BioPAX-Explorer ^4^ is a versatile Python package dedicated to the development of BioPAX data analysis scripts and software applications. BioPAX-Explorer can be installed using pip. A Docker image is proposed to deploy it easily with Jupyter and the Fuseki triple store ^5^.

## 4. Results

We outline the main functionalities integrated into the BioPAX-Explorer package and provide a brief illustration of their usage.

### 4.1 BioPAX-Explorer as a modular library

Several modules are built on top of Core BioPAX-Explorer, among them:

- a BioPAX data model, coded in Python, with all relations between classes;
- a Pattern oriented domain specific language (DSL), to query BioPAX datasets without writing SPARQL, that includes a SPARQL query generator and a query module using RDFLib;
- an Object to Triple Mapper, that converts any SPARQL query results to a list of BioPAX entities, as instances of the BioPAX Python classes;
- a graph integration module to seamlessly convert a query result in a graph data structure; the graph can then be easily analysed using reference graph analysis Python packages, NetworkX and Graph-tool.

### 4.2 BioPAX-Explorer to formulate queries

#### 4.2.1 Using existing Patterns

BioPAX-Explorer offers directly usable patterns, representing canonical queries. So far 40 patterns have been coded and are available, whose results will be similar to those provided by some of the Paxtools patterns used as reference. The patterns can be directly reused to extract data, without the necessity to study the BioPAX model in details.

The Python pattern definitions are available in the “Rack” sub package. For flexibility, data awareness and learning, the internal generated SPARQL queries can be displayed with the method

**`querylist=PatternExecutor().queries(p)`** (see supplementary data). These queries can subsequently be repurposed in alternative contexts. As a use case, Figure 4 shows the usage of the pattern **`inComplexWith`**, that extracts all occurrences of complexes that are made of at least 2 proteins. In the “Rack” pattern collection, we also coded original patterns. Some of them can be used to detect elements which are not compliant with the BioPAX specification. For example **`notBlackboxComplexInComplex`**, which could have been used in Juigné *et al*. (2023) instead of having to manually design the low-level SPARQL query.

**Fig 4.**
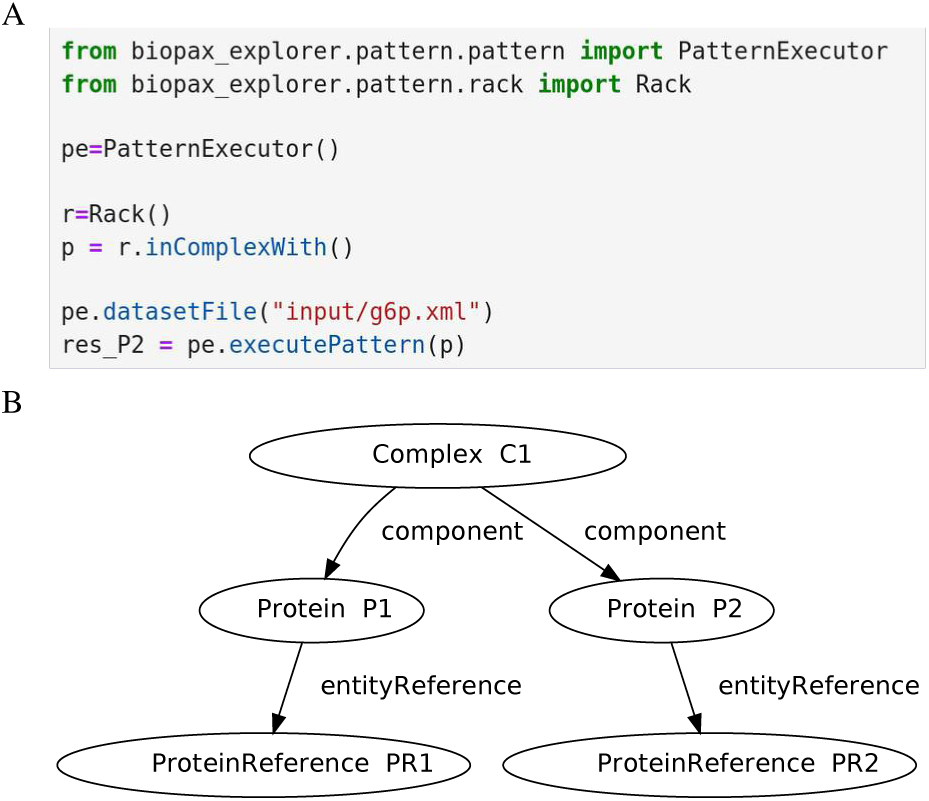
A Query Pattern available in the canonical Pattern collection, provided by BioPAX-Explorer. This query pattern is designed for retrieving molecular complexes and the protein references of the constituent proteins. The listing (A) shows a Python code snippet used to reuse the pattern. The diagram (B) represents the resulting query pattern executed on a dataset from a local file using “executePattern”. The results can be accessed through the variable “res_p2” (A), which is structured as a list of rows. Each row comprises a list of entities, with each entity corresponding to a node in the diagram.

#### 4.2.2 Creating new patterns

BioPAX-Explorer also allows users to create their own custom extraction patterns in Python, in an object-oriented syntax, by directly manipulating the classes in the BioPAX model. The “Pattern” syntax is inspired from Paxtools Patterns. It is expressive and object-oriented. Figure 5 illustrates this functionality, by showing Python code for crafting a Pattern and its execution. The result comprises class instances, which correspond to BioPAX entities, with attributes populated with corresponding values. Their types include primitive types or classes, conforming precisely to the BioPAX specifications.

**Fig 5.**
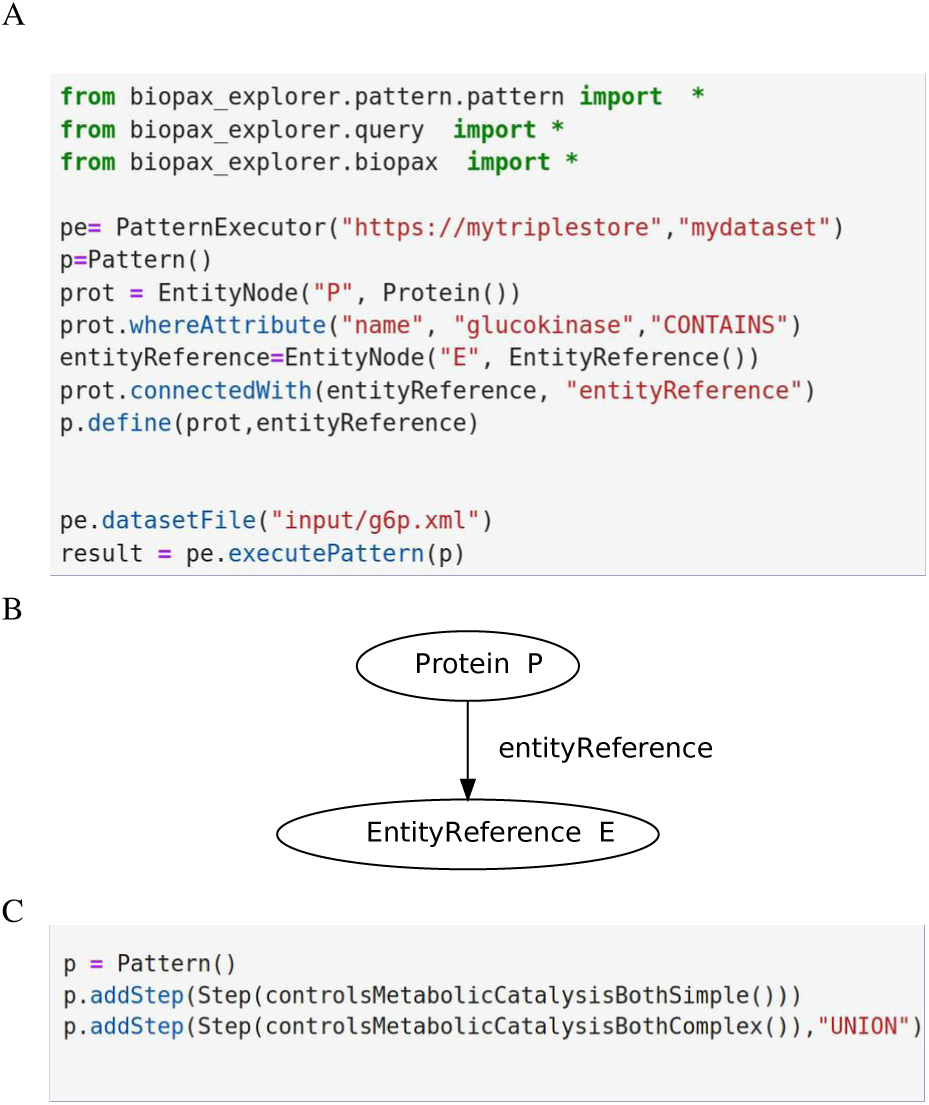
A Query Pattern, created using the BioPAX-Explorer object oriented Pattern syntax, the embedded BioPAX query DSL (Domain Specific Language). Listing (A) shows the Python code used to create the Pattern from scratch. Diagram (B) is a visual representation of the pattern that can be executed on a dataset, from a triple store accessible on the network. This pattern defines any protein with an entity reference associated with it via the “entityref” relationship in accordance with the BioPAX specification. A help module can be used to discover the properties of any classes in the BioPAX model and make it easier to write such a Pattern. The listing (C) shows an advanced Pattern construction using sub Patterns and Set operators.

Complex patterns can be designed by combining other patterns, considered as steps (Figure 5-C). Each step can be either a simple pattern, an operation to combine patterns (intersection, union…), or filter instances. The “Rack” module uses these mechanisms.

#### 4.2.3 Executing query

BioPAX-Explorer includes a query generator that operates like a compiler, transforming the DSL pattern into a graph representation, and subsequently into SPARQL. The SPARQL query is then sent to the execution engine. The patterns, internally converted to SPARQL, are executed, and the outcomes are mapped as a list of instances of classes of the Python BioPAX model.

### 4.3 BioPAX-Explorer to explore large datasets as graphs

A query pattern execution results in a list containing the instances representing BioPAX entities, as described in Figure 6-A. A Pattern can be executed to extract all attributes and relations of each entity (Figure 6-B).

**Fig 6.**
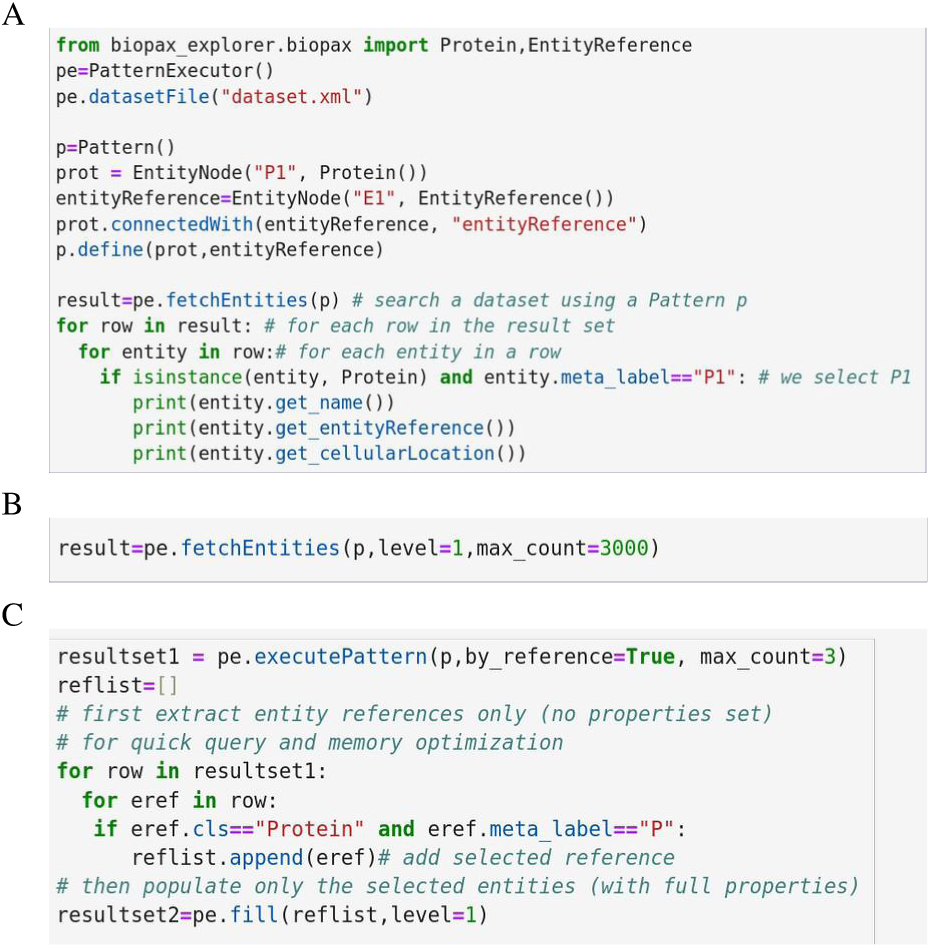
Listing A shows how the result of a pattern execution can be processed. The results are class instances corresponding to BioPAX entities, populated with all attributes values, available in a 2 dimensions list. Users can choose to extract the full populated entities (listing B) or only references first and the full entities in a second time (listing C). The level parameter defines the extent of the neighbourhood (1 for only selected entities,2 for direct relation, 3 relations at one more level of distance). Variable “pe” have previously been defined.

Different features allow to explore large datasets:

1) Selecting references on the entities, which include only their identifier and class name, and then, for the selected instances, populating all the attributes (Figure 6-C). The “level” parameter automatically includes entities from an extended or narrow neighborhood. This enables progressive exploration of very large knowledge graphs. For example, if a pattern defines an interaction node, all controlled entities associated with the interaction instances found in the dataset will be populated on demand and made available in Python via the methods **`interaction.get\_controlled()`**. By this way we can easily explore the neighbourhoods close to the entities we are focused on, without retrieving the full set of network interactions.
2) Traversing the relations of the chosen entities, as outlined in the BioPAX specification. It is noteworthy that certain attribute values, treated as relations, are themselves instances of BioPAX classes. For instance, in a **Complex** entity, there is an attribute called “component” that refers to a list of **`PhysicalEntities`**. Each of these entities **(if it is a `Protein`)**, is referencing an **`EntityReference`**, thus creating a cascading structure. As a result, this exploration traces a path within the knowledge graph.
3) Entities can also be accessed as nodes in a graph by automatically converting a list of entities into a graph data structure. In this context, each node is a BioPAX entity and each edge is a relationship between entities. We obtain a data structure equivalent to a property graph as showed in Figure 7. A “labeled property graph” is a data model where nodes represent entities with associated key-value pairs, while edges denote relationships between nodes, also supporting properties. One of the advantages of representing data in the form of a graph data structure in memory is to facilitate post-processing. It supports complex operations such as path-search, that are useful when processing pathways but are beyond the scope of SPARQL. In BioPAX-Explorer, we provide two in memory graph backend implementations, NetworkX for ease of use and Graph-tool for its performances on large graphs. These characteristics facilitate the rapid development of analysis algorithms while maintaining a reproducible workflow from large datasets.

**Fig 7.**
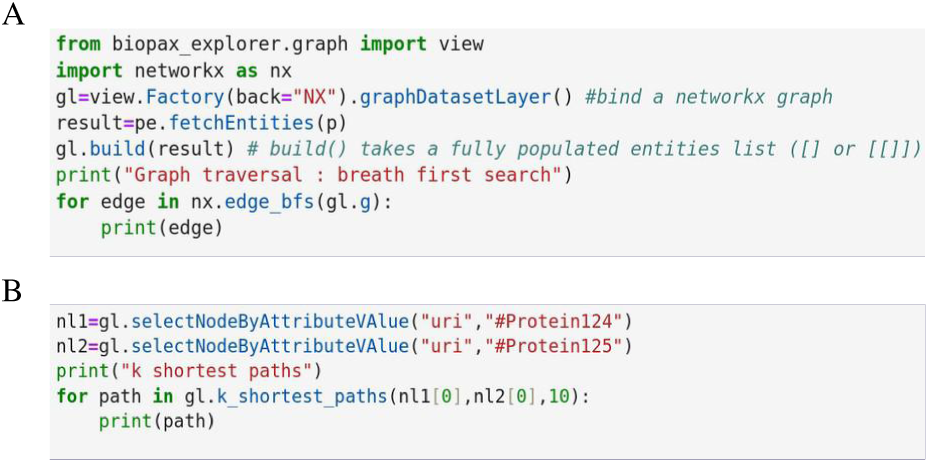
Graph mapping: an simplified graph mapper is provided to convert a result set directly to a graph data structure. Then classical traversal (A) and path search algorithm (B)can be used. All attributes of the entities are available as node properties.

### 4.4 Functionalities in Graph visualization

BioPAX-Explorer includes several functionalities for data visualization. The data, transformed into a property graph, can be easily visualized using NetworkX. Users can also export the graph as GRAPHML, making it compatible with standalone software such as Cytoscape (Shannon *et al*., 2003) or GEXF in Gephi (Bastian *et al*., 2009). For seamless integration with Jupyter notebooks, BioPAX-Explorer incorporates a graph viewer (Franz *et al*., 2015) (see supplementary data).

### 4.5 Data Exchange and Validation

Originally designed for BioPAX dataset exploration, BioPAX-Explorer also offers robust object-oriented modeling capabilities that extend its applications beyond exploration. Particularly, it can be used to generate BioPAX data sources. This functionality involves the **BPSerializer** class, which streamlines the conversion of lists of BioPAX entities into RDF/XML BioPAX files. By employing a straightforward object-oriented syntax, developers can instantiate entities using constructors and setters, enabling the rapid assembly of complex datasets.

Additionally, BioPAX-Explorer provides a versatile validation mechanism through a dedicated module. Unlike predefined validation processes, this module allows to construct validation methods tailored to specific requirements. For instance, users can list all entities whose URIs are not provided or those that are not unique or all proteins lacking cellular localization information. This approach ensures flexibility and adaptability to different validation scenarios encountered in practical applications (see supplementary data).

### 4.6 Use case of BioPAX-Explorer

As a use case of BioPAX-Explorer, we conducted a comparative analysis of two datasets from Reactome (Milacic *et al*., 2024), namely the “event -*Homo sapiens*” and “event -*Mus musculus*” datasets, both adhering to BioPAX level 3 standards. Figure 8 depicts the results of this analysis. The simple queries (A) can easily be designed as Patterns. More advanced queries (B) would have required a detailed understanding of the model; however, we reused BioPAX-Explorer prebuild Patterns to accelerate their implementation. The script is available in the supplementary material as Jupyter notebooks.

**Fig 8.**
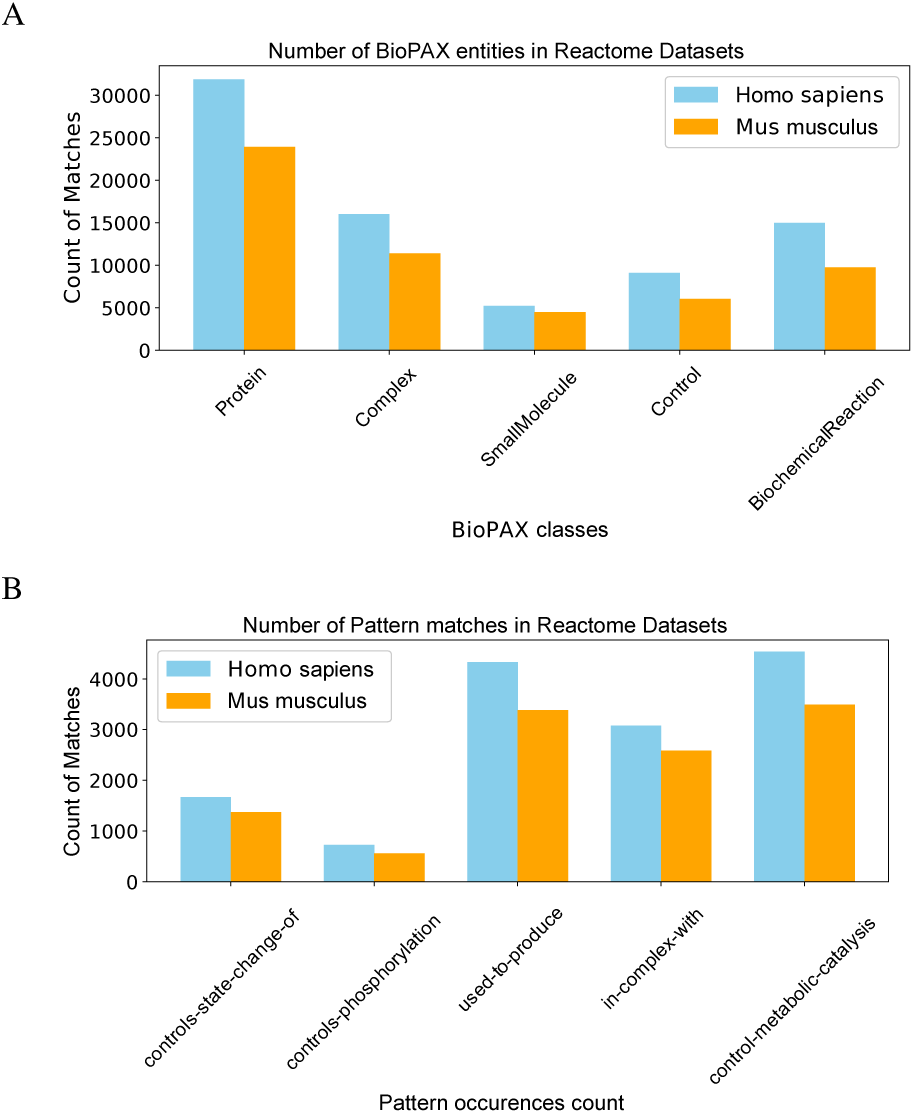
Datasets comparison between species. We utilized user-defined BioPAX-Explorer patterns to conduct a straightforward metrics analysis (A) on Reactome’s BioPAX datasets. Furthermore, a more sophisticated analysis (B) was also conducted by using some predefined patterns available within the Rack module.

### 4.7 Comparison with Paxtools

To evaluate the reliability of the pattern-based query system within the BioPAX-Explorer package, we selected a set of patterns from the “Rack” prebuilt collection that closely resemble existing Paxtools patterns.

We utilized BioPAX files provided by Reactome for the species *Mus musculus* and *Homo sapiens*. These files were integrated into a triple store, and we queried the datasets using the patterns within BioPAX-Explorer.

In parallel, we queried the datasets using semantically equivalent Paxtools patterns directly on the files.

Our analysis revealed that the number of matches obtained from both systems, BioPAX-Explorer and Paxtools, were of the same order of magnitude (refer to supplementary data online).

## 5. Discussion and perspectives

BioPAX-Explorer simplifies the utilization and integration of Biological Pathway Exchange (BioPAX) data for Python users. The package was developed to address the complexity of BioPAX datasets, while keeping the power of RDF and SPARQL thanks to a simple object-oriented syntax. It offers a fully BioPAX-compatible object model, featuring advanced query capabilities, model reconstruction, and validation.

BioPAX-Explorer facilitates exhaustive exploration of large datasets, bypassing memory limits with SPARQL query generation and a memory cache mechanism. It accelerates the development of Python-based biological network analysis software, fostering transparent and reproducible workflows based on the BioPAX OWL standard.

By generating internally the SPARQL queries, it enables easy data extraction, making it accessible to a wide audience of Python users.

BioPAX-Explorer enables to:

- explore exhaustively BioPAX dataset from files or SPARQL endpoints, even for very large dataset;
- use Entity oriented simple queries and result sets construction without the necessity to learn a complex query language like SPARQL;
- learn how to write complex SPARQL queries for RDF datasets dedicated to biological networks, that can be reused within external tools;
- accelerate the development of Python based biological network analysis software by allowing the generation of a graph data structure from a query on BioPAX dataset;
- facilitate the creation of a transparent and reproductible workflow, from data to analysis, based on BioPAX OWL standard, from start to end.

The finding of the comparison with Paxtools underscores the robustness and reliability of the BioPAX-Explorer package’s pattern-based query system in comparison to established tools. However, interpreting the results of this comparison remains challenging. The prebuilt patterns provided in BioPAX-Explorer do not have direct equivalents in Paxtools. There are inherent differences in semantics and design between the two systems, and establishing systematic equivalences would necessitate expert knowledge in both environments.

Additionally, from all patterns of BioPAX-Explorer, it remains always possible to examine the SPARQL code that was automatically executed by BioPAX-Explorer. The analysis of the SPARQL allows to verify that the queries executed faithfully represent the intended semantics. By this way, we ensure that the results are transparent and explainable. This verification process operates independently of the specific tools employed and maintains a direct correlation with the underlying data. Such a mechanism not only enhances the reliability of the results but also fosters confidence in the reproducibility of the data analysis process and enhance the explainability of the obtained results.

Data analysis reproducibility ensures that the results of an analysis can be consistently and reliably obtained by others using the same data and methodology. BioPAX-Explorer, that uses internal SPARQL can contribute to reproducibility and explicability in several ways. The use of internal SPARQL promotes reproducibility through standardized querying. The implementation of query traceability enhances explicability, which refers to the clarity and transparency of the query process, allowing users to easily understand and interpret how the requested information is retrieved and derived.

The SPARQL queries generated by BioPAX-Explorer can be reused in different contexts, independently of BioPAX-Explorer. This enables users to leverage the queries for other purposes or within alternative tools, providing flexibility and facilitating the creation of diverse applications or workflows.

Moreover, BioPAX-Explorer can be used for the exchange of BioPAX datasets while ensuring their integrity and compliance with domain-specific standards. This approach enhances the efficiency and reliability of data management processes. On the whole, the ability of BioPAX-Explorer to facilitate the editing and validation of datasets should help to increase the usage of BioPAX.

Another advantage is the ability to leverage online BioPAX data sources made available as SPARQL endpoints, a standardized and remote-accessible interface for querying RDF data. In this context, we believe that BioPAX-Explorer can become an essential part of a distributed and standardised infrastructure for making biological interaction network data collectively available.

Future prospects for BioPAX-Explorer involve establishing models associations to connectable data sources, such as ChEBI or UniProt. It would be beneficial to explore the generalization of the mapping approach between ontologies, RDF and Object-Oriented implementations. This could streamline the integration of internal data from research teams with publicly available datasets, enhancing collaborative research efforts. Additionally, there is an emphasis on integrating this approach into intelligent visualization and exploration tools that are generic and applicable to various types of OWL ontologies, exploiting the versatility of Semantic Web technologies.

## Acknowledgements

We acknowledge the GenOuest bioinformatics core facility (https://www.genouest.org) for providing the computing infrastructure.

## 7. Supplementary data

Supplementary data are available at https://fjrmoreews.github.io/biopax-explorer/

## Footnotes

1 https://github.com/pwin/owlready2

2 https://github.com/RDFLib/rdflib

3 https://github.com/pallets/jinja/

4 https://fjrmoreews.github.io/biopax-explorer

5 https://forgemia.inra.fr/pegase/biopax-explorer#installation

